# Preservation and storage effects on river sediment microbiomes: Implications for community stability and ecological inference

**DOI:** 10.64898/2026.09.05.749588

**Authors:** Joeselle M. Serrana, Malte Posselt

## Abstract

Sample preservation and storage can alter microbial communities between sampling and analysis, influencing the interpretation of environmental microbiome data. We investigated how preservation method, storage temperature, and storage duration affect river sediment microbiomes using 16S and 18S rRNA gene amplicon sequencing. Sediments were preserved in ethanol, nucleic acid preservation (NAP) buffer, or without a preservative, and stored at room temperature or frozen at −20°C or −80°C for up to eight weeks. We report that preservation method, storage conditions, and their interaction influenced estimates of microbial diversity and community composition. Differences between treatments were detectable after approximately 18 h and generally increased with storage time. Frozen samples remained closest to their treatment-specific baseline communities, whereas room-temperature storage, particularly without a preservative, produced the largest changes. Notably, ethanol and NAP buffer preservation reduced, but did not prevent, changes during room-temperature storage. Preservation also influenced taxonomic patterns and the outcome of phylogenetic null-model analyses. Frozen storage generally retained the sample’s assembly-metric estimates, with stronger signatures of deterministic community assembly, whereas room-temperature storage shifted the inferred balance toward stochastic processes. Our results show that preservation and storage conditions can affect both the microbial community measured and the ecological conclusions drawn from it. Among the conditions tested, frozen storage provided the best preservation of sediment microbial communities and should be preferred when samples cannot be processed immediately.

**Graphical Abstract:** 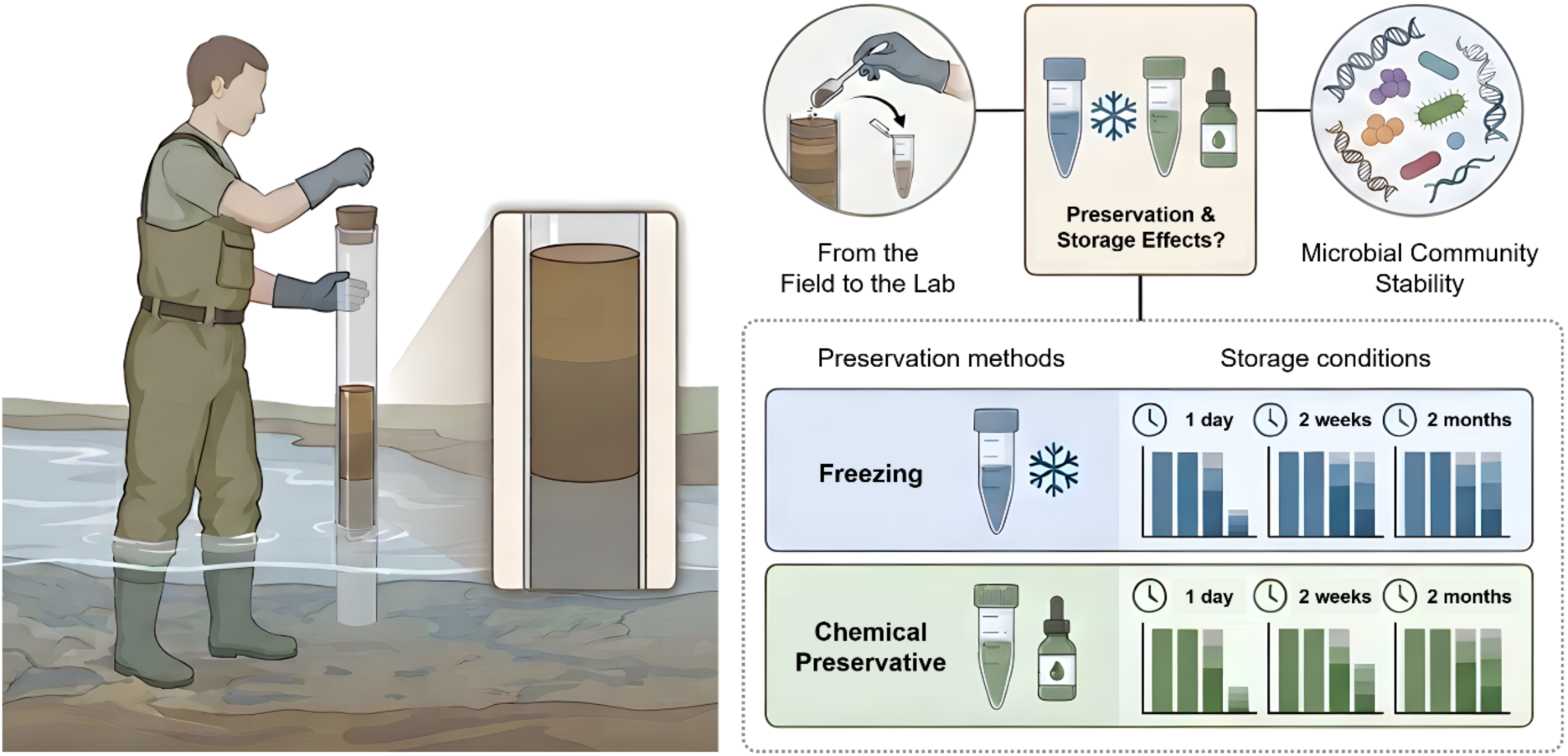

## Introduction

Accurate characterization of environmental microbiomes depends on preserving the microbial community present at the time of sampling (Tennant et al., 2022; Warren et al., 2024). However, sample handling, storage conditions, and the time between collection and nucleic acid extraction can alter the community that is ultimately measured (e.g., Ahn et al., 2022; Serrana & Watanabe, 2022; Tennant et al., 2022). In environmental matrices, e.g., river sediments, changes during storage can promote the growth of opportunistic taxa, accelerate nucleic acid degradation, and alter community composition, obscuring the original ecological and biogeochemical signals present in the sample (Guerrieri et al., 2021; Brasell et al., 2022; Keneally et al., 2024; Warren et al., 2024; Guerrieri et al., 2026). Hence, maintaining microbiome integrity from field collection to laboratory processing remains a challenge across amplicon sequencing, metagenomics, and other omics-based approaches (Pawlowski et al., 2022).

Standardized sampling protocols help reduce methodological variability and improve comparability among studies (Guerrieri et al., 2021; Bensch et al., 2022; Guerrieri et al., 2026). Immediate subsampling and field preservation, either by flash-freezing or by rapidly applying chemical preservatives, is commonly recommended to limit post-collection changes caused by oxygen exposure, temperature fluctuations, and repeated freeze-thaw cycles (Pribyl et al., 2021; Wu et al., 2021; Bensch et al., 2022; Wietz et al., 2022; Trivedi et al., 2022; Stica et al., 2026). These measures aim to retain microbial diversity and community composition as close as possible to their state at collection, reducing bias in subsequent ecological interpretation (Pawlowski et al., 2022; Keneally et al., 2024).

Chemical preservatives have become particularly important because they allow samples to be stored and transported at ambient temperatures, reducing the logistical challenges and costs of maintaining a continuous cold chain from field collection to laboratory processing (e.g., Song et al., 2016; Dully et al., 2021; Bizzozzero et al., 2023). Common preservatives include Longmire’s buffer, RNAlater, DNA/RNA Shield, ethanol-based solutions, and other reagents designed to stabilize nucleic acids when immediate freezing is impractical (Pawlowski et al., 2022; Guerrieri et al., 2026). One example is the nucleic acid preservation (NAP) buffer, which contains EDTA, sodium citrate tribasic dihydrate, and ammonium sulfate and has been shown to maintain the integrity of both DNA and RNA at room temperature (Camacho-Sanchez et al., 2013; Menke et al., 2017). Despite their increasing use, the effectiveness of these preservation approaches in maintaining sediment-associated microbiomes during prolonged storage remains poorly understood, particularly compared with conventional freezing-based methods.

An additional challenge is that preservation effects may vary across microbial groups. Previous studies have reported substantial differences in nucleic acid stability among prokaryotic and eukaryotic microorganisms during preservation, storage, and processing (e.g., Brauer & Bengtsson, 2022; Alan et al., 2025; Ashley-Wheeler et al., 2026). Preservation treatments may unequally affect different components of the microbiome, introducing taxon-specific biases that influence estimates of diversity, community composition, and ecological relationships. Such biases affect interpretations of ecosystem functioning, microbial interactions, and assembly processes. Systematic evaluations of preservation and storage conditions are therefore needed to determine how different components of sediment microbiomes respond over time and to identify approaches that best maintain microbiome stability.

In this study, we investigated how preservation method, storage temperature, and storage duration affect the stability of river sediment microbiomes using 16S and 18S rRNA gene amplicon sequencing. Specifically, we compared microbiomes preserved with 95% ethanol, NAP buffer, or no preservative and stored at room temperature (RT), −20°C, and −80°C for up to 8 weeks. We then evaluated changes in microbial diversity, community composition, stability, and community assembly processes across preservation and storage conditions. We hypothesized that samples maintained at low temperatures throughout transport and storage would exhibit greater microbiome stability, reflected in lower community divergence from baseline communities over time.

By assessing responses across prokaryotic and microbial eukaryotic communities, this study offers practical guidance for preserving sediment microbiomes and advances the development of standardized protocols for freshwater sediment sampling, storage, and processing. Additionally, the findings will help improve the reliability of environmental nucleic acid approaches for freshwater biodiversity assessment, ecological monitoring, and long-term environmental surveillance.

## Methods

### Experimental design

Surface sediment samples were taken from the River Fyris (Uppsala, Sweden) by collecting the upper 5 cm of the benthic sediments with a sediment corer. The collected sediment cores were immediately homogenized and subsampled in the field into sterile 15-mL collection tubes, with sediment subsamples filling approximately 5 mL of volume in each tube. In total, 18 subsamples were prepared and subjected to different field preservation and laboratory storage conditions. Three subsamples were transported from the field by keeping them in a cooler with dry ice and stored at −80°C in the laboratory, while two sets of three subsamples were preserved in a 1:2 sediment-to-preservative volume ratio (10 mL) of 95% ethanol (EtOH) and in nucleic acid preservation (NAP) buffer (microbial DNA-free; Sigma-Aldrich, Steinheim, Germany). Nine samples were prepared without a field preservative and transported at ambient temperature; 1 set was stored in the laboratory at room temperature (∼22°C; RT), and 2 sets were placed in −20°C and −80°C freezers upon arrival (∼2 hrs. after sample collection). All samples were then sampled at five time points: ∼18 h in storage after field collection, and after 2, 4, 6, and 8 weeks (**Supplementary Table S1**). Samples processed at ∼18 h after field collection were used as the baseline reference to evaluate microbiome stability during storage. For each preservation treatment, three replicate storage tubes were then repeatedly subsampled throughout the storage-duration experiment.

### Amplicon sequencing and data processing

Total genomic DNA was extracted from ∼300 mg of sediment from each sample using the DNeasy PowerSoil Pro kit (Qiagen GmbH, Hilden, Germany), following the manufacturer’s instructions with minor modifications to the cell lysis time and elution steps. We measured the quantity and quality of the extracted DNA using the NanoPhotometer N60 (IMPLEN GmbH, Germany). A blank sample served as the negative extraction control. Aliquots of the extracted DNA were sent to Novogene Europe (Cambridge, UK) for amplicon sequencing using two barcode regions: (1) the prokaryotic V4-V5 region of the 16S rRNA gene amplified with the 515F-907R primers, and (2) the eukaryotic V4 region of the 18S rRNA gene amplified using the 528F-706R primers. All PCR reactions contained 10 ng of DNA template, 2 µM of forward and reverse primers, and 15 µL of Phusion® High-Fidelity PCR Master Mix (New England Biolabs), and were amplified under the following conditions: initial denaturation for 1 min at 98°C, followed by 30 cycles of denaturation at 98°C for 10 s, annealing at 50°C for 30 s, elongation at 72°C for 30 s, and a final extension step of 5 min at 72°C. Quality-checked PCR products were pooled at equal concentrations and purified using a Qiagen Gel Extraction Kit (Qiagen, Germany). Following the manufacturer’s instructions, sequencing libraries were prepared using the TruSeq DNA PCR-Free Sample Preparation Kit (Illumina, USA). The quality of the resulting amplicon library was assessed on the Qubit 2.0 Fluorometer (Thermo Scientific) and the Agilent Bioanalyzer 2100 system. Qualified amplicon libraries were then sequenced on an Illumina NovaSeq 6000 platform (Illumina, San Diego, CA, USA), generating 250 bp paired-end reads.

The raw reads were checked for quality using FastQC v0.11.9 (Andrews, 2010). The paired-end reads were demultiplexed for each sample based on their unique indices, then truncated to remove the barcode and primer sequences. The demultiplexed paired-end reads were then quality-screened and processed via the DADA2 v1.4.0 pipeline (Callahan et al., 2016). The rarefaction curves calculated for ACE, Chao1, Shannon, and observed ASV indices approached a plateau or a stable point (**Supplementary Figure S1**). Read filtering was performed using the following quality parameters: a maximum expected error of 2, and reads were truncated at lengths of 220 bp for the forward reads and 200 bp for the reverse reads. Reads matching against the PhiX genome were also filtered out. Amplicon sequence variants (ASVs) were inferred by denoising the quality-filtered reads with the DADA2 error model. The reads were then dereplicated, merged, and sequences containing chimeric reads were removed (**Supplementary Table S2**). For each ASV, the taxonomic assignment was performed using the *assignTaxonomy* function in DADA2, implementing the RDP naive Bayesian classifier method (Wang et al., 2007) against the Silva v138.2 (Quast et al., 2013) database for 16S rRNA and the Protist Ribosomal Reference (PR2) v5.0.0 (Guillou et al., 2012) database for 18S rRNA data with the minimum bootstrapping support set to default. Multiple sequence alignments of the prokaryotic and eukaryotic datasets were performed using DECIPHER v2.26.0 (Wright, 2015). A phylogenetic tree was then constructed with FastTree v2.1.11 (Price et al., 2009) using the GTR nucleotide substitution model.

Contaminant ASVs were identified by their higher abundance in the negative control than in the samples using decontam v1.24 (Davis et al., 2018) and were subsequently removed from the dataset. Moreover, ASVs with < 10 counts were excluded (Amir et al., 2017). For the 16S rRNA data, only ASVs assigned to bacteria and archaea were retained, while ASVs assigned to chloroplasts or mitochondria were discarded. For the 18S rRNA data, ASVs annotated as bacteria, archaea, or embryophytes were filtered out. Sample sequence abundances were normalized to the median sequencing depth across samples by scaling ASV counts according to differences in library size prior to downstream analyses. The resulting ASV tables (**Supplementary Tables S3** and **S4**) and phylogenetic trees for each dataset were compiled into phyloseq objects using the phyloseq v1.48.0 package (McMurdie & Holmes, 2013) and microtables using the microeco v2.0.0 package (Liu et al., 2026) for downstream analyses.

### Visualization and data analyses

All statistical analyses were conducted in R v4.6.1 using the microeco v2.3.0 package (Liu et al., 2026) and associated packages for ecological, phylogenetic, and statistical analyses. Prokaryotic (16S rRNA gene) and microbial eukaryotic (18S rRNA gene) datasets were analyzed separately.

#### Estimating alpha diversity and community composition

Alpha diversity was quantified using observed richness, Shannon diversity, and Pielou’s evenness. The effects of preservation treatment, storage duration, and their interaction on alpha diversity were evaluated separately for the 16S and 18S datasets using linear mixed-effects models, with tube identity included as a random effect to account for repeated measurements through time. Microbial community composition was evaluated using Bray-Curtis dissimilarities calculated from ASV abundance data. Differences in community composition among preservation treatments, storage durations, and their interaction were assessed using permutational multivariate analysis of variance (PERMANOVA; 999 permutations) implemented in the vegan package (Oksanen et al., 2015). Principal coordinate analysis (PCoA) was used to visualize patterns in community composition. To assess whether significant PERMANOVA results were associated with differences in within-group variability, homogeneity of multivariate dispersions was evaluated using PERMDISP with 999 permutations. To determine whether preservation treatments differed shortly after collection, we also compared microbial community composition among the 18 h baseline samples using PERMANOVA and PERMDISP analyses based on Bray-Curtis dissimilarities.

#### Community stability and taxonomic response analyses

Community stability was quantified as the Bray-Curtis dissimilarity between each stored sample and the corresponding baseline community sampled after 18 h within the same preservation treatment. Because preservation treatments already differed after 18 h (see Results), stability analyses were performed relative to treatment-specific baseline communities rather than a common baseline across treatments. For each preservation method, we calculated a baseline centroid as the mean community composition of the 18 h replicates, then calculated Bray-Curtis dissimilarities between each stored sample and this baseline centroid. Community divergence was defined as:

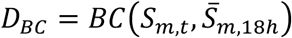

where *D_BC_* is the Bray-Curtis distance-to-baseline, *S_m_*_,*t*_ is the community composition of preservation treatment *m* at storage duration *t*, and *̅S_m_*_,18ℎ_ is the centroid (mean community composition) of the 18 h baseline replicates for the same preservation treatment. Lower values indicate greater stability, whereas larger values indicate greater divergence from treatment-specific baseline conditions.

Taxonomic responses to preservation treatment and storage duration were evaluated using relative abundance data aggregated at the phylum level for prokaryotic communities and division level for eukaryotic communities. Taxa that showed consistent increases or decreases in relative abundance over time were interpreted as indicators of preservation-related biases and storage-induced community shifts.

#### Inference of community assembly processes

Community assembly processes were evaluated using phylogenetic null-model analyses of the *trans_nullmodel* class functions implemented in the microeco v2.3.0 package (Liu et al., 2026). For each dataset, we calculated the β-nearest taxon index (βNTI) using randomized null communities. βNTI values were used to quantify deviations in phylogenetic turnover from null expectations and infer the relative importance of deterministic assembly processes (Liu et al., 2017). βNTI values > +2 and < −2 were interpreted as evidence of variable selection and homogeneous selection, respectively. For pairwise comparisons where |βNTI| < 2, Raup-Crick dissimilarities based on Bray-Curtis distances (RCBray) were calculated to further distinguish stochastic assembly mechanisms. Comparisons with RCBray > 0.95, RCBray < −0.95, and |RCBray| ≤ 0.95 were classified as dispersal limitation, homogenizing dispersal, and undominated processes (including ecological drift), respectively (Stegen et al., 2013). The relative contributions of homogeneous selection, variable selection, dispersal limitation, homogenizing dispersal, and ecological drift were then quantified both across the complete datasets and separately for each preservation treatment to evaluate whether preservation methods influenced inferred community assembly patterns.

## Results

### Amplicon sequencing statistics and overall microbial community structure

Amplicon sequencing generated 5,231,025 and 5,137,145 raw reads for the 16S and 18S datasets, respectively. Following quality filtering, denoising, and read merging, 34.1% and 87.1% of reads were retained as merged sequences in the 16S and 18S datasets, respectively. Subsequent chimera removal and contaminant screening retained 93.0% and 98.4% of merged reads for downstream analyses (**Supplementary Table S2**). After processing, sequencing depth ranged from approximately 14,000 to 50,000 reads per sample for the 16S dataset and from 45,000 to 71,000 reads per sample for the 18S dataset. After normalization, the final datasets comprised 10,050 prokaryotic and 8,616 eukaryotic amplicon sequence variants (ASVs).

Rarefaction curves approached or nearly asymptotes for all samples, indicating sufficient sequencing depth to capture the majority of prokaryotic and eukaryotic diversity present in the sediment samples (**Figure S1**). The rarefaction curves also showed clear differences in how preservation methods affected 16S diversity, while 18S curves remained comparatively clustered. For 16S, ethanol and NAP buffer at room temperature, along with immediate −80°C storage, produced consistently higher richness across metrics. The no-preservative RT treatment also showed elevated observed ASV richness relative to the other treatments. In contrast, the −20°C no-preservative condition displayed uniformly lower richness and Shannon values. For 18S, all preservation methods produced tightly overlapping curves for richness and Shannon.

Across all samples, prokaryotic communities were dominated by Pseudomonadota (28.6%), Chloroflexota (13.9%), Actinomycetota (10.6%), Bacillota (9.5%), and Acidobacteriota (7.5%), accounting for approximately 70% of the total prokaryotic community. Additional abundant groups included Bacteroidota (5.6%) and Thermodesulfobacteriota (5.5%) (**Figure 2A**). Eukaryotic communities were largely composed of Opisthokonta (49.7%), Alveolata (15.9%), Chlorophyta (10.2%), Rhizaria (9.6%), and Stramenopiles (9.2%), which represented more than 94% of the total eukaryotic community (**Figure 2B**).

**Figure 1.**
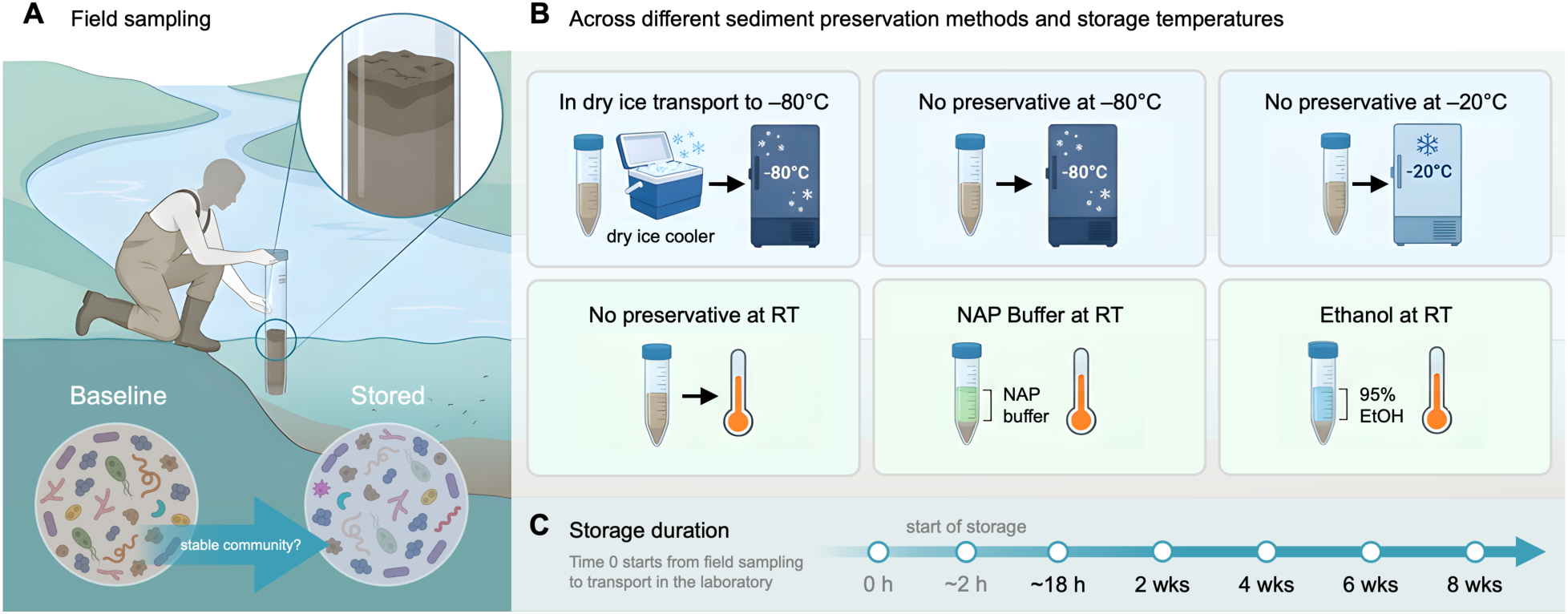
Experimental design of the study. (A) Surface sediment (0-5 cm) from the River Fyris (Uppsala, Sweden) was collected using a sediment corer, homogenized on site, and subsampled into aliquots. (B) Subsamples were assigned to preservation treatments, i.e., 95% ethanol (EtOH), nucleic acid preservation (NAP) buffer at room temperature (RT), or no preservative, and stored at different temperature regimes (i.e., RT, −20°C, or −80°C). One treatment was transported on dry ice and stored at −80°C. (C) Samples were then processed after ∼18 h and after 2, 4, 6, and 8 weeks of storage. Microbial community responses (i.e., diversity, community composition, and stability) were characterized from the prokaryotic and microbial eukaryotic profiles generated using 16S and 18S rRNA gene amplicon sequencing. The graphics used in this figure were created using FigureLabs (https://figurelabs.com).

**Figure 2.**
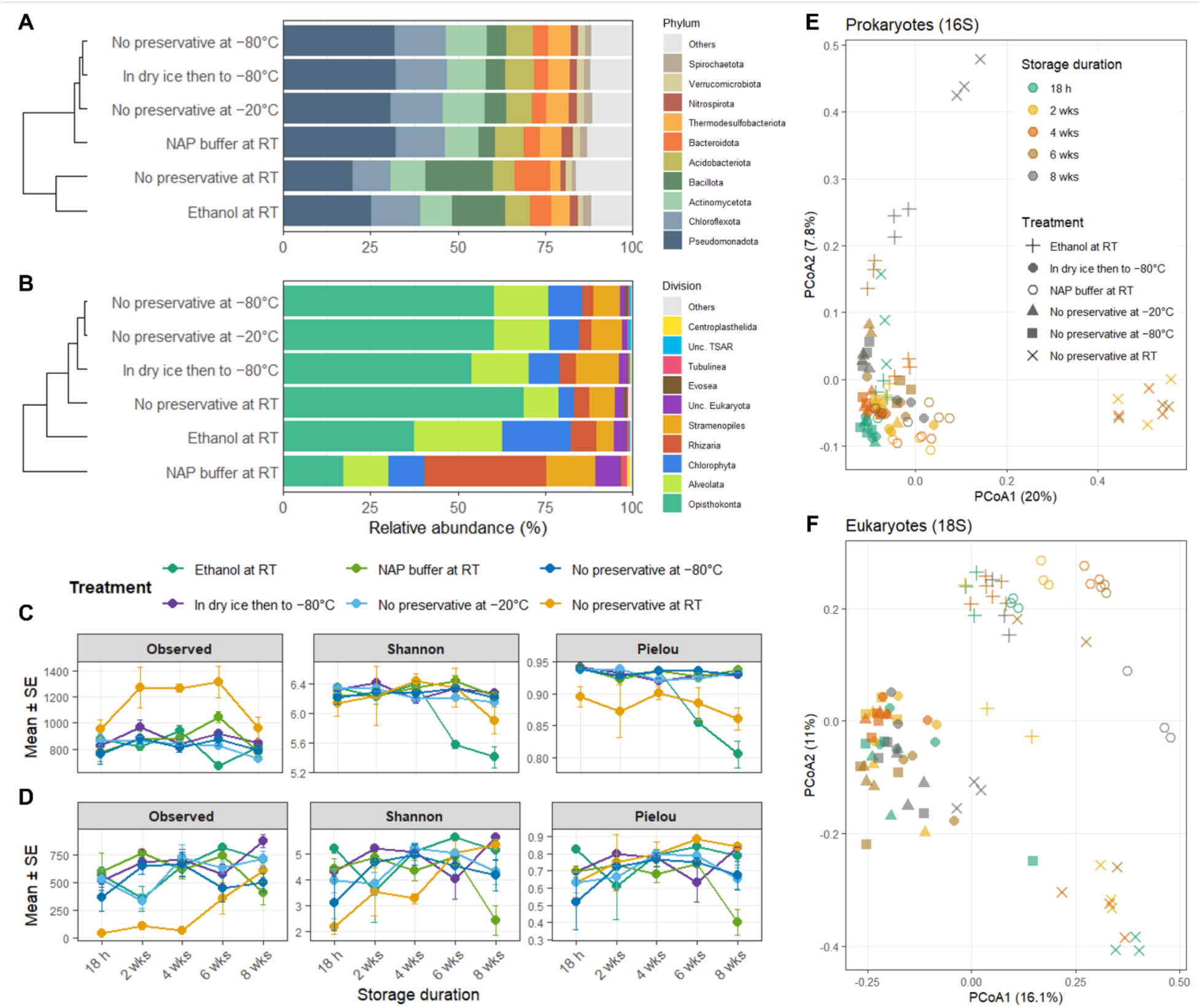
Effects of preservation method and storage duration on prokaryotic (16S) and eukaryotic (18S) microbial community diversity and composition. Community composition across preservation treatments for 16S (A) and 18S (B) datasets. Relative-abundance profiles of the top 10 taxa are shown with dendrograms summarizing similarity among treatments. Temporal changes in alpha diversity of the prokaryotic (C) and eukaryotic (D) communities, with observed richness, Shannon diversity, and Pielou’s evenness measured from samples preserved using different storage treatments and analyzed after 18 h, 2, 4, 6, and 8 weeks of storage. Points represent treatment means, and error bars indicate ± standard error (SE) of the replicates. Principal coordinate analysis (PCoA) of Bray-Curtis dissimilarities for prokaryotic (E) and eukaryotic (F) communities. Point colors represent storage duration (18 h, 2, 4, 6, and 8 weeks), and point shapes denote preservation treatments.

Notably, frozen treatments grouped closely together, with −80°C storage and dry-ice transport to −80°C showing highly similar community profiles, and −20°C positioned nearby but slightly shifted, indicating modest compositional change relative to fully frozen storage. All RT treatments (ethanol, NAP, and no-preservative) grouped together and clearly apart from the frozen set, reflecting a shared shift in community structure under warm storage regardless of preservative chemistry. Moreover, community composition among baseline samples processed approximately 18 h after collection already showed treatment-specific differences, indicating that preservation-associated effects were detectable shortly after sample collection (**Supplementary Figure S2**). Hence, the baseline communities were assessed as treatment-specific references to evaluate storage-induced changes over time.

### Preservation method and storage duration alter microbial diversity and community composition

Preservation treatment and storage duration significantly influenced microbial alpha diversity, as indicated by linear mixed-effects models that accounted for repeated measurements within storage tubes (**Figure 2C-D** and **Supplementary Table S5**). In the prokaryotic dataset, both preservation treatment and storage duration significantly influenced the observed richness, Shannon diversity, and Pielou’s evenness (*P* < 0.001 in all cases). Similarly, preservation treatment and treatment × time interactions significantly influenced all diversity metrics (*P* < 0.033) in the eukaryotic dataset, while storage duration significantly affected observed richness and Shannon diversity (P < 0.007) but not Pielou’s evenness (*P* = 0.053). Overall, alpha diversity trajectories were generally more stable under frozen storage conditions, whereas unpreserved RT samples showed greater temporal variation and larger deviations from baseline values.

Moreover, the preservation treatment and storage duration significantly influenced microbial community composition. Principal coordinate analyses revealed clear separation among preservation treatments and storage durations in both datasets (**Figure 2E-F**). Temporal progression through storage was associated with increasingly divergent community compositions, although the magnitude and direction of change differed among preservation strategies. Preservation and storage condition explained 62.3% and 60.7% of the total variation in prokaryotic and eukaryotic community composition, respectively. For 16S, preservation treatment explained 22.2% of the variation in Bray-Curtis dissimilarities (PERMANOVA, *R^2^*= 0.222, *P* = 0.001), whereas storage duration accounted for 10.7% of the variation (*R^2^* = 0.107, *P* = 0.001) (**Supplementary Table S6**). The interaction between preservation treatment and storage duration explained an additional 29.4% of the variation (*R^2^* = 0.294, *P* = 0.001), indicating that temporal changes in community composition depended strongly on the preservation strategy employed.

Similar patterns were observed in the 18S dataset, where preservation treatment explained 29.5% of the variation in community composition (*R^2^* = 0.295, *P* = 0.001), whereas storage duration explained 5.9% (*R^2^* = 0.059, *P* = 0.001). The interaction between treatment and storage duration accounted for a further 25.3% of the variation (*R^2^* = 0.253, *P* = 0.001), demonstrating that preservation effects varied across storage durations. PERMDISP analyses also indicated significant differences in multivariate dispersion among preservation treatments in both the 16S dataset (*F* = 134.86, *P* = 0.001) and the 18S dataset (*F* = 15.90, *P* = 0.001), suggesting that preservation/storage condition effects influenced not only community composition but also within-treatment variability.

### Frozen storage maintains the highest microbial community stability across preservation treatments

Preservation treatments and storage conditions significantly influenced microbial community composition after 18 h in both datasets, explaining 45.3% and 58.9% of the variation in 16S and 18S communities, respectively (PERMANOVA, *P* = 0.001; **Supplementary Table S7**). Since treatment-associated differences were already evident at baseline, subsequent stability analyses quantified community divergence relative to treatment-specific 18 h baselines. For 16S, preservation treatment (*F_5,59_* = 267.01, *P* < 0.001), storage duration (*F_4,59_* = 338.87, *P* < 0.001), and their interaction (*F_20,59_* = 18.16, *P* < 0.001) strongly influenced community stability. The fitted model explained 98.1% of the total variation in Bray-Curtis distance-to-baseline. Similarly, community stability in the 18S dataset was significantly affected by preservation treatment (*F_5,60_* = 59.26, *P* < 0.001), storage duration (*F_4,60_* = 75.48, *P* < 0.001), and the treatment × time interaction (*F_20,60_* = 4.50, *P* < 0.001), with the model explaining 92.0% of the variation in community divergence.

Across both datasets, Bray-Curtis distances increased progressively with storage time, indicating divergence from the baseline community composition (**Figure 3A-B**). However, divergence magnitude differed markedly among preservation treatments. Samples stored without preservative at RT consistently showed the greatest divergence from baseline communities and differed significantly from all other treatments from 2 weeks onward in both datasets (Tukey HSD, *P* < 0.001 in most comparisons).

**Figure 3.**
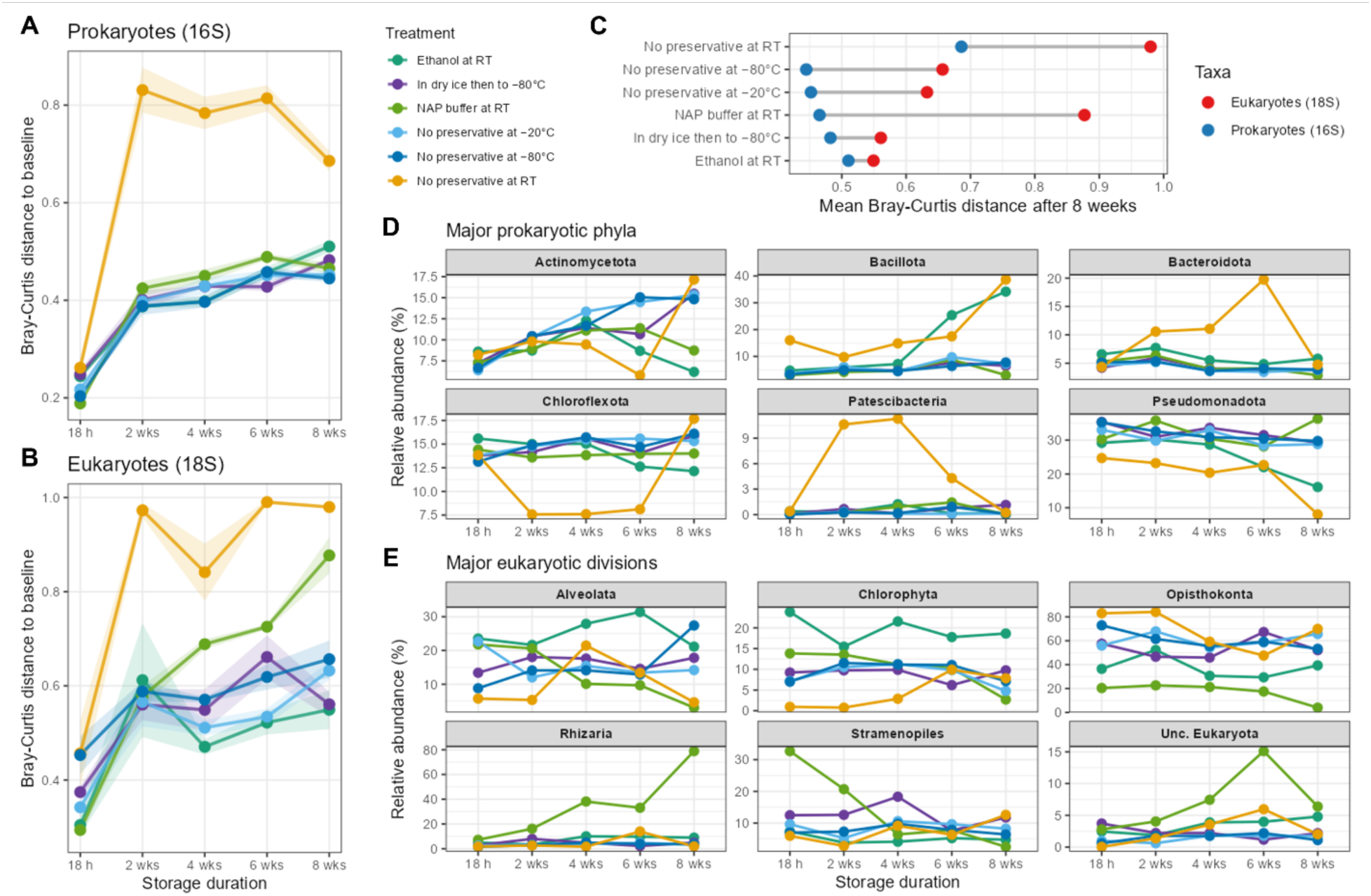
Community stability across preservation treatments and storage durations. Bray-Curtis distance-to-baseline for (A) prokaryotic and (B) eukaryotic communities. Lines represent mean values across replicates and shaded ribbons indicate ± SE. Increasing Bray-Curtis distances indicate progressive divergence from baseline community composition during storage. (C) Mean Bray-Curtis distance-to-baseline of the microbial communities after 8 weeks of storage. Lower values indicate greater community stability and preservation effectiveness. Statistical analyses of the effects of preservation treatment, storage duration, and their interaction on community stability are presented in **Supplementary Table S7**. Temporal trajectories of the top six major prokaryotic phyla (D) and eukaryotic divisions (E) across preservation treatments and storage durations. Lines represent mean relative abundances of three replicates. Taxa were selected based on the highest variance in relative abundance across all treatment and storage-duration combinations. Broad taxonomic patterns for the 10 most abundant taxa per dataset are presented in **Supplementary Figure S2**.

Relative community stability (mean Bray-Curtis distance-to-baseline) at 8 weeks further highlighted substantial differences among preservation methods (**Figure 3C** and **Supplementary Table S7**). For prokaryotic communities, storage at −80°C without preservative showed the greatest stability (mean Bray-Curtis distance = 0.445), followed by storage at −20°C (0.452), NAP buffer at RT (0.465), dry ice followed by −80°C storage (0.482), and ethanol preservation (0.510). As expected, the no-preservative RT-stored treatment showed the greatest divergence from baseline communities (0.686) and was the least stable. Eukaryotic communities showed a similar overall pattern, although treatment rankings differed among preservation methods. Ethanol preservation exhibited the lowest mean Bray-Curtis distance after 8 weeks (0.549), followed closely by dry ice transport at −80°C storage (0.561). However, freezing was the most consistently effective preservation strategy across both marker datasets, whereas the performance of chemical preservatives was marker-dependent. In contrast, NAP buffer (0.877) and no-preservative RT storage (0.980) showed substantially greater divergence from baseline communities, indicating that NAP preservation was markedly less effective for eukaryotic communities than for prokaryotes. These results show that microbial communities become increasingly dissimilar to their baseline composition during storage, but freezing consistently kept communities closer to baseline than RT storage with preservative.

### Preservation and storage condition effects are unevenly distributed across microbial lineages

Broad taxonomic patterns varied among preservation treatments and storage durations for both prokaryotic and eukaryotic communities (**Supplementary Figure S2**). Preservation-associated shifts were particularly evident in several dominant bacterial phyla and eukaryotic divisions, indicating that storage effects were not uniformly distributed across microbial groups. Temporal trajectories of the six most abundant prokaryotic phyla revealed treatment-specific responses through time (**Figure 3D**). Several dominant phyla, including Pseudomonadota, Bacillota, and Bacteroidota, exhibited pronounced shifts under RT storage, whereas frozen treatments generally maintained abundance profiles closer to those observed at baseline conditions.

In particular, the no-preservative RT treatment showed the largest deviations from baseline abundance patterns across multiple prokaryotic phyla. Similar trends were observed for eukaryotic communities (**Figure 3E**), where major divisions, i.e., Alveolata, Chlorophyta, Opisthokonta, Rhizaria, and Stramenopiles, responded differently to preservation treatment and storage duration. Several eukaryotic divisions exhibited pronounced temporal fluctuations under RT storage conditions, whereas frozen treatments generally retained more stable taxonomic profiles. Broad taxonomic responses were further evident in heatmaps of the 10 most abundant bacterial phyla and eukaryotic divisions (**Supplementary Figure S2**), indicating that storage-associated community divergence reflected widespread, coordinated shifts across the microbiome rather than changes restricted to a few dominant taxa. These patterns indicate that preservation effects act unevenly across the microbiome, altering the relative abundance of specific lineages and potentially influencing downstream ecological interpretation. The observed shifts further suggest that preservation-associated biases are taxon-specific, leading to systematic changes across multiple microbial lineages that may influence ecological conclusions when preservation procedures are not standardized.

### Inferred community assembly processes differ among preservation and storage conditions

Phylogenetic null-model analyses revealed that inferred community assembly patterns from one sediment sample varied among preservation treatments and between microbial groups. βNTI distributions were predominantly negative across preservation treatments, indicating that phylogenetic turnover was generally lower than expected under null-model expectations and suggesting a strong influence of deterministic processes on community structure (**Figure 4A-B**). However, the magnitude and variability of βNTI values differed among preservation treatments, particularly in the eukaryotic dataset. Overall assembly- process summaries indicated that prokaryotic communities were predominantly shaped by homogeneous selection, whereas eukaryotic communities showed a substantially greater contribution from ecological drift (**Figure 4C** and **Supplementary Table S8**).

**Figure 4.**
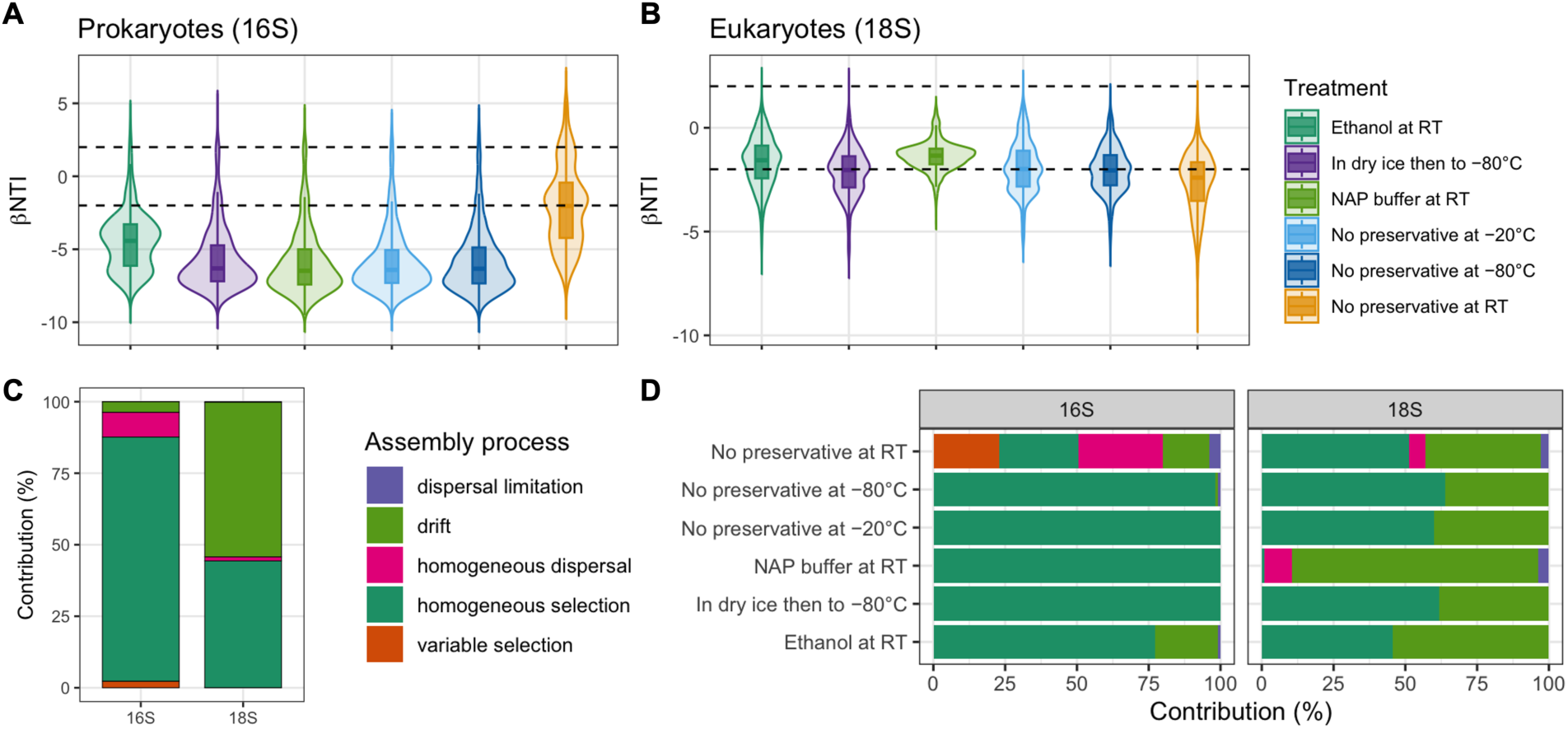
Effects of preservation treatments on inferred patterns of microbial community assembly. Distribution of β-nearest taxon index (βNTI) values for (A) prokaryotic (16S) and (B) eukaryotic (18S) communities across preservation treatments. Dashed horizontal lines indicate the βNTI thresholds (±2) commonly used to distinguish deterministic from stochastic assembly processes. (C) Overall relative contributions of variable selection, homogeneous selection, dispersal limitation, homogeneous dispersal, and ecological drift inferred from βNTI and RCBray null-model analyses. (D) Treatment-specific assembly-process contributions for prokaryotic (16S) and eukaryotic (18S) communities. Values represent the percentage of pairwise community comparisons assigned to each assembly process.

Treatment-specific analyses, however, revealed considerable variation among preservation methods (**Figure 4D)**. In prokaryotic communities, frozen storage treatments were dominated by homogeneous selection, accounting for 98.1-100% of pairwise comparisons. NAP-preserved samples stored at RT similarly exhibited complete dominance of homogeneous selection (100%). In contrast, ethanol-preserved samples showed a greater contribution from ecological drift (21.9%), while unpreserved samples stored at RT showed the most pronounced shift in assembly patterns, with homogeneous selection accounting for only 27.6% of pairwise comparisons. Under these conditions, homogenizing dispersal (29.5%), variable selection (22.9%), ecological drift (16.2%), and dispersal limitation (3.8%) all contributed substantially to inferred community assembly.

In contrast, eukaryotic communities responded differently to preservation treatment. Frozen storage treatments remained primarily associated with homogeneous selection, accounting for 60.0-63.8% of pairwise comparisons, although ecological drift contributed 36.2-40.0%. Ethanol-preserved samples exhibited a greater contribution of drift (54.3%) than homogeneous selection (45.7%). The largest departure from deterministic assembly occurred in NAP-preserved samples stored at room temperature, where ecological drift accounted for 85.7% of pairwise comparisons and homogeneous selection contributed less than 1%. No-preservation RT samples also showed substantial stochastic influence, with drift accounting for 40.0% of inferred assembly processes.

Overall, frozen storage consistently retained stronger signatures of deterministic assembly, whereas RT treatments, particularly in eukaryotic communities, increased the relative importance of stochastic processes. These findings suggest that preservation and storage conditions may affect not only estimates of microbial diversity and community composition but also ecological interpretations derived from community assembly analyses.

## Discussion

Understanding how preservation influences microbiome data is essential as environmental nucleic acid approaches become increasingly integrated into freshwater monitoring and ecological assessment (Keneally et al., 2024; Allan et al., 2025; Zou et al., 2025). In this study, we evaluated how preservation method, storage temperature, and storage duration influence river sediment microbiomes, with particular emphasis on microbial diversity, community composition, temporal stability, and the ecological patterns inferred from phylogenetic null-model analyses. By comparing freezing-based and chemical preservation strategies across prokaryotic and eukaryotic communities, we aimed to assess how sample handling influences microbiome characterization and downstream ecological interpretation.

### Microbiome stability depends strongly on preservation and storage conditions

Microbial diversity and community composition changed progressively during storage, but the magnitude of these changes differed substantially among preservation strategies. Notably, we already observed preservation-associated effects after 18 h, indicating that sample handling can influence microbiome composition shortly after collection. This finding suggests that methodological biases may arise much earlier than often assumed, and that variation introduced during transport and short-term storage may substantially contribute to observed differences in the microbiome among studies (Tennant et al., 2022; Maghini et al., 2024). Hence, standardization of field-to-laboratory workflows may be as important as the choice of long-term preservation strategy itself.

We highlight that frozen microbial communities remained most similar to their treatment-specific baseline communities, whereas prolonged room-temperature storage led to increasing divergence over time (**Figure 3A-C**). In particular, the samples stored at −80°C showed the least divergence from baseline communities and retained community compositions most similar to their treatment-specific baseline reference communities. Even −20°C storage generally outperformed room temperature storage, indicating that lowering the temperature effectively limits biological and chemical processes that continue after sample collection. Given that microbial growth, enzymatic activity, DNA degradation, and cell lysis can alter community profiles during storage (Jenkins et al., 2018; Mason-Jones et al., 2022; Smenderovac et al., 2024; Warren et al., 2024; Ashley- Wheeler et al., 2026), the superior performance of frozen storage is therefore consistent with the expectation that low temperatures suppress post-collection biological activity and retain community compositions closest to their treatment-specific baseline reference communities.

Similar benefits of frozen preservation have been reported across a variety of environmental matrices, including soils, freshwater samples, and marine sediments, supporting the continued recommendation to maintain a cold chain whenever logistically feasible (Romanazzi et al., 2015; Pulido Barriga et al., 2025; Stica et al., 2026). Moreover, we observed that preservation effects were not limited to shifts in community composition. Significant differences in multivariate dispersion among treatments indicate that some preservation strategies also altered the consistency of replicate community profiles, increasing among-replicate variability and potentially reducing the reproducibility of microbiome measurements.

On the other hand, both ethanol and NAP buffer preservation also reduced community drift compared with unpreserved samples stored at room temperature, demonstrating their utility for field-based studies under logistical constraints. However, neither preservative completely prevented compositional changes during prolonged storage. These findings are consistent with recent sediment-based evaluations showing that preservation strategies influence biodiversity estimates and community composition across multiple taxonomic groups, although the magnitude and direction of preservation effects can vary among habitats, markers, and extraction workflows (Guerrieri et al., 2026). This suggests that chemical preservatives should be viewed as mitigating, rather than eliminating, storage artifacts. Their effectiveness likely depends on storage duration, sample matrix characteristics, and taxonomic composition (Tatangelo et al., 2014; Song et al., 2016; Sales et al., 2019; Dully et al., 2021). In particular, sediment samples pose preservation challenges because microorganisms occupy heterogeneous, particle- associated microhabitats that can limit preservative penetration and potentially sustain localized biological activity (Torti et al., 2018; Dumoulin et al., 2026; Paul et al., 2026). Still, chemical preservatives may provide practical alternatives when immediate freezing of sediment samples is not feasible.

### Preservation effects differ between microbial groups

Prokaryotic and eukaryotic communities responded differently to preservation and storage conditions. Several dominant eukaryotic divisions showed pronounced temporal fluctuations during room temperature storage, whereas many bacterial phyla remained comparatively stable (**Figure 3D-E**). These differences indicate that preservation bias is unlikely to act uniformly across the microbiome and suggest that protocols optimized for bacterial communities may not perform equivalently for microbial eukaryotes. This has important implications for contemporary environmental DNA and microbiome studies, which increasingly characterize multiple domains of life simultaneously to better understand ecosystem structure and function (Afonso et al., 2024; Sandré et al., 2025; Serrana & Watanabe, 2025; Serrana et al., 2026). Differential preservation responses among microbial groups may affect estimates of biodiversity, community composition, and ecological interactions, particularly in studies integrating bacterial, protistan, fungal, and metazoan assemblages (Guerrieri et al., 2026).

Additionally, we observed that taxonomic shifts further demonstrate that preservation effects are not uniform across microbial communities (**Figure 3D-E; Supplementary Figure S2**), with some taxa remaining relatively stable regardless of storage conditions, whereas others showed marked increases or decreases in abundance over time. These patterns likely reflect differences in growth potential during storage, nucleic acid degradation rates, resistance to preservation treatments, or DNA persistence after cell death (Song et al., 2016; Wietz et al., 2022; Ashley-Wheeler et al., 2026). These taxon-specific responses matter because many biomonitoring frameworks rely on indicator taxa or changes in community composition as evidence of ecological condition (Astudillo-García et al., 2019; Simonin et al., 2019; Gómez-Repollés et al., 2026; Serrana et al., 2026). Storage-induced shifts in the abundance of sensitive lineages may therefore introduce biases that could be misinterpreted as biological responses to environmental stressors (Meyer et al., 2026). This is consistent with broader evidence that differences in preservation performance can alter the relative representation of taxa in sediment samples, potentially biasing metabarcoding-based assessments of community composition. Preservation-associated taxonomic biases have been reported across multiple studies, highlighting the importance of standardized sample handling (e.g., Dully et al., 2021; Brasell et al., 2022; Mejbel et al., 2022; Paul et al., 2026). Thus, comparisons among studies using different preservation strategies should be interpreted with caution, particularly when evaluating the abundance of specific microbial lineages.

### Preservation artifacts propagate into ecological inference

Preservation effects extended beyond diversity and taxonomic composition to the patterns inferred from phylogenetic null-model analyses. (**Figure 4C-D**). In prokaryotic communities, frozen storage treatments and NAP-preserved samples remained dominated by homogeneous selection, whereas ethanol-preserved and unpreserved room temperature samples exhibited substantially greater contributions from stochastic and dispersal-related processes. In eukaryotic communities, preservation effects were even more pronounced, with NAP-preserved samples showing a marked increase in ecological drift and a corresponding reduction in homogeneous selection. However, because preservation treatments also differed in community dispersion and diversity, assembly-process estimates should be interpreted as inferred patterns rather than direct measurements of ecological mechanisms. Importantly, we show that the same original sediment can yield markedly different conclusions from community-assembly analyses depending on how the sample was preserved.

The predominance of homogeneous selection in frozen samples suggests that freezing at −20°C or −80°C more effectively maintained the ecological structure of their treatment-specific reference communities. Ecologically, homogeneous selection typically indicates that similar environmental filters act consistently across communities, favoring taxa with comparable ecological characteristics and producing predictable community composition (Stegen et al., 2013). The strong homogeneous-selection signal observed in frozen samples therefore suggests that these preservation approaches better retained community organization present in their treatment-specific baseline communities during storage. In contrast, storage treatments associated with increased ecological drift imply a growing influence of stochastic changes in community composition. Such patterns may arise when random gains, losses, or fluctuations in taxa occur during preservation and storage, reducing community predictability and increasing variation among samples (Stegen et al., 2013; Fodelianakis et al., 2021).

Particularly notable was the shift observed in unpreserved room temperature samples and, for eukaryotic communities, in NAP-preserved samples stored at room temperature (**Figure 4D**). In these treatments, deterministic assembly signals weakened while stochastic processes became increasingly important. If preservation artifacts increase the apparent importance of drift, microbial communities may appear more stochastically structured than they actually are under natural conditions. Conversely, preservation treatments that preserve strong signatures of homogeneous selection may better retain the ecological relationships among taxa, suggesting that different preservation strategies can produce fundamentally different ecological interpretations from the same environmental sample. Hence, preservation should not be viewed solely as a technical consideration. Instead, it can be a source of variation that propagates from microbiome characterization to downstream ecological interpretation and hypothesis testing.

Differences in ecological process estimates highlight a frequently overlooked consequence of sample preservation. Preservation effects are commonly evaluated in terms of diversity loss or compositional shifts, yet methodological variation may also propagate into higher-order ecological interpretations. As microbial ecology increasingly relies on phylogenetic null models to infer mechanisms, e.g., environmental filtering, dispersal dynamics, and ecological drift (Choudoir et al., 2022; Wang et al., 2023; Flores et al., 2025), preservation-induced changes in community structure may influence conclusions regarding the processes governing biological communities. This issue may be particularly important for environmental DNA and biomonitoring applications that characterize multiple taxonomic groups simultaneously (Afonso et al., 2024; Sandré et al., 2025). If preservation methods differ in their tendency to identify deterministic or stochastic assembly signals, differences among datasets may reflect methodological variation rather than genuine ecological patterns.

Preservation effects should therefore be considered alongside other sources of uncertainty, including sampling design, sequencing protocols, and bioinformatic processing, when interpreting microbial community assembly patterns. Nonetheless, our findings reinforce the growing evidence that post-collection handling can substantially influence microbiome characterization and downstream ecological interpretation (Jenkins et al., 2018; Sales et al., 2019; Brasell et al., 2022; Warren et al., 2024).

### Implications for environmental monitoring

We demonstrate that preservation and storage practices can substantially influence estimates of microbial diversity, community composition, and inferred ecological processes, highlighting the importance of accounting for methodological artifacts in environmental microbiome studies. As environmental nucleic acid approaches become increasingly integrated into long-term monitoring programs, biodiversity assessments, ecosystem restoration initiatives, and aquatic biosecurity frameworks (Taberlet et al., 2012; Littlefair et al., 2022), distinguishing genuine ecological change from preservation-induced variation becomes increasingly critical. The reliability of these applications ultimately depends on the accurate preservation of molecular signals from sample collection through laboratory analysis. The rapid emergence of preservation-associated effects observed in this study indicates that methodological biases may be introduced shortly after collection and can continue to accumulate during storage. Accordingly, differences among preservation workflows may influence both microbial taxon detection and the ecological conclusions drawn from community data. Such effects are particularly important for environmental monitoring programs that seek to identify temporal trends, detect ecological disturbances, or compare datasets generated across multiple studies and institutions (Hartig et al., 2024).

Overall, immediate freezing provided the most reliable preservation of river sediment microbiomes, whereas ethanol and nucleic acid preservation (NAP) buffer offered practical alternatives when maintaining a continuous cold chain was impractical. However, the differing responses among microbial groups, along with the influence of storage on inferred community assembly processes, show that preservation effects extend beyond measures of diversity and composition alone. Unstandardized sample handling may therefore reduce comparability among studies, obscure biological signals, and complicate the interpretation of long-term ecological change. Establishing standardized preservation workflows is therefore essential for improving the reproducibility, comparability, and ecological reliability of freshwater microbiome and environmental nucleic acid research (Bunholi et al., 2023; Shea et al., 2023; Yang et al., 2023). Moreover, our results emphasize that sample preservation should be regarded as a critical component of study design and quality assurance in biomonitoring and environmental surveillance programs.

While our results provide valuable insights into how preservation and storage conditions influence sediment microbiomes, we still note several limitations when interpreting these findings. The study was conducted using sediments from a single river system, and preservation responses may differ among environmental matrices, ecosystem types, and climatic regions (Lever et al., 2015; van der Heyde et al., 2020; Mejbel et al., 2022). Thus, the relative performance of preservation methods and the magnitude of storage-induced effects may not be universally transferable across habitats.

Additionally, the differences between prokaryotic and eukaryotic responses should be interpreted cautiously, given that 16S and 18S datasets are derived from different marker genes with distinct amplification efficiencies, copy-number variation, and taxonomic resolution. Finally, the analyses were based on DNA sequence data and therefore cannot distinguish between changes in active microbial communities and changes arising from nucleic acid degradation, extracellular DNA persistence, or the accumulation of DNA from inactive organisms (Zöhrer et al., 2026). Future studies spanning multiple habitats, environmental conditions, preservation protocols, and molecular targets, including complementary RNA-based approaches, will help determine the generality of the patterns observed here and further refine best-practice recommendations for environmental microbiome research.

## Conclusion

Preservation methods and storage conditions strongly influenced the diversity, composition, stability, and inferred assembly processes of river sediment microbiomes. Frozen storage most effectively maintained communities that resembled treatment-specific baseline conditions. Although ethanol and nucleic acid preservation (NAP) buffer provided practical alternatives when maintaining a continuous cold chain was not feasible, neither completely prevented storage-induced changes. Hence, preservation artifacts influenced not only microbiome characterization but also the ecological inferences derived from environmental nucleic acid datasets. Moreover, preservation methods altered the inferred balance between deterministic and stochastic assembly processes, demonstrating that sample handling can influence ecological interpretation as well as community composition. These findings highlight the importance of considering sample preservation as a potential source of bias in freshwater microbiome studies and support the adoption of standardized preservation workflows for ecological monitoring and biodiversity assessment.

## Author Contributions

**Joeselle M. Serrana:** Conceptualization; Methodology; Investigation; Formal Analysis; Data Curation; Visualization; Validation; Writing – Original Draft. **Malte Posselt:** Conceptualization; Validation; Resources; Supervision; Project Administration; Funding Acquisition; Writing – Review & Editing.

## Acknowledgments

We thank Arild Gustafsson and Run Tian for their assistance in the fieldwork and sample transport. The authors also thank Elias Broman for feedback on the study’s initial experimental design. Open-access funding is provided by Stockholm University. The study was supported by the Swedish Research Council for Sustainable Development (FORMAS) (Grant Number 2021-02059). J.M.S. was supported by the Stockholm University Center for Circular and Sustainable Systems (SUCCeSS) (Project No. 30002687) postdoc funding. The computations were performed on the supercomputer Dardel at PDC in KTH Royal Institute of Technology, with access and resources provided by the National Academic Infrastructure for Super-computing in Sweden (NAISS) through projects NAISS 2025/22-1105, 2025/23-467, and 2026/4-1362. The graphical abstract and figure graphics were created using FigureLabs (https://figurelabs.com).

## Conflicts of Interest

The authors declare no conflicts of interest.

## Data Availability Statement

Raw sequence data are available at the NCBI SRA repository under BioProject ID PRJNA1336108. Additional data from the analyses presented in this paper are available in the Supplementary Material. The corresponding visualization, analysis input data, and codes are also deposited on GitHub: https://github.com/jserrana/preservation-mtb.

## Notes

https://github.com/jserrana/preservation-mtb

### Competing Interest Statement

The authors have declared no competing interest.

