## Supplementary Figures for "Preservation and storage effects on river sediment microbiomes: Implications for community stability and ecological inference"

**Authors & Affiliations**

Joeselle M. Serrana[1,2,*] and Malte Posselt[2]

[1]Stockholm University Center for Circular and Sustainable Systems (SUCCeSS), Stockholm University, 106 91 Stockholm, Sweden
[2]Department of Environmental Science (ACES), Stockholm University, 106 91 Stockholm, Sweden

### Supplementary Table Legends

**Supplementary Table S1.** Metadata for all samples included in the preservation experiment. Information includes sample identifiers, preservation treatments, storage conditions, and storage durations.

**Supplementary Table S2.** Sequencing and bioinformatic processing statistics for the 16S and 18S rRNA gene datasets. Values represent read counts retained after each processing step, including quality filtering, denoising, merging, chimera removal, and final non-chimeric sequence recovery.

**Supplementary Table S3.** Amplicon sequence variant (ASV) abundance table for the prokaryotic (16S rRNA gene) dataset. Rows represent ASVs and columns represent samples. Taxonomic assignments are provided from kingdom to species level based on the SILVA v138.2 database.

**Supplementary Table S4.** Amplicon sequence variant (ASV) abundance table for the eukaryotic (18S rRNA gene) dataset. Rows represent ASVs and columns represent samples. Taxonomic assignments are provided from domain to species level based on the PR2 v5.1.0 database.

**Supplementary Table S5.** Results of linear mixed-effects models evaluating the effects of preservation treatment, storage duration, and their interaction on observed richness, Shannon diversity, and Pielou's evenness in the 16S and 18S datasets. Pairwise comparisons among treatments and storage durations were performed using Tukey's honestly significant difference (HSD) tests.

**Supplementary Table S6.** Results of Bray-Curtis-based community composition analyses for the 16S and 18S datasets. The table includes PERMANOVA results for preservation treatment, storage duration, and their interaction, together with homogeneity-of-dispersion (PERMDISP) analyses and principal coordinate analysis (PCoA) statistics.

**Supplementary Table S7.** Statistical analyses of microbial community stability quantified as Bray-Curtis distance-to-baseline. Results include ANOVA, post hoc comparisons, divergence-rate models, and preservation-treatment rankings after 8 weeks of storage. Lower Bray-Curtis values indicate greater similarity to treatment-specific baseline communities and therefore greater preservation effectiveness.

**Supplementary Table S8.** Relative contributions (%) of variable selection, homogeneous selection, dispersal limitation, homogenizing dispersal, and ecological drift inferred from βNTI and RCBray null-model analyses. Assembly processes were quantified for the complete 16S and 18S datasets and separately for each preservation treatment to evaluate the effects of sample preservation on inferred microbial community assembly patterns.

### Supplementary Figures

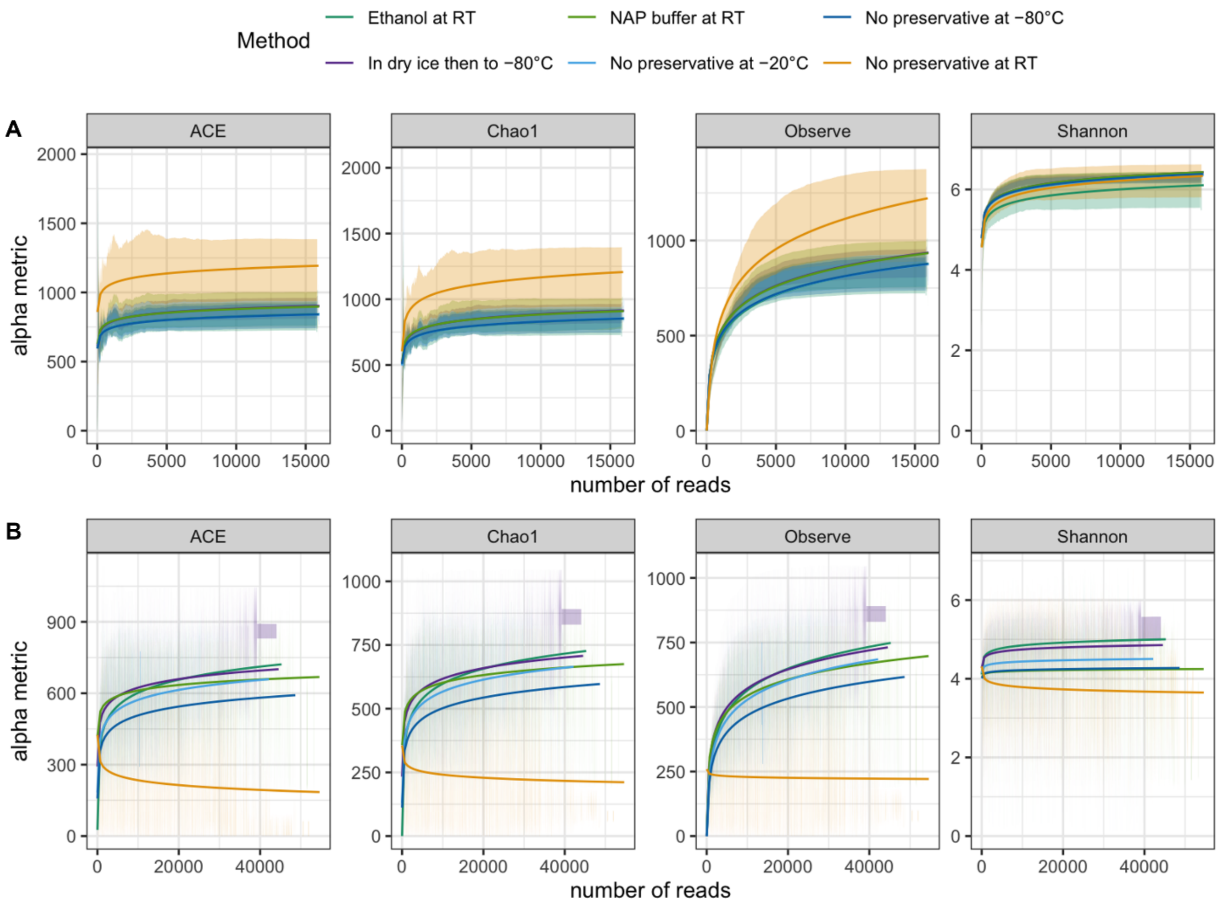


**Supplementary Figure S1.** Rarefaction curves of the amplicon sequence variant (ASVs) reads. (A) 16S and (B) 18S dataset showing ACE, Chao1, Observed ASVs, and Shannon diversity across increasing read depth. Shaded regions indicate replicate variability.

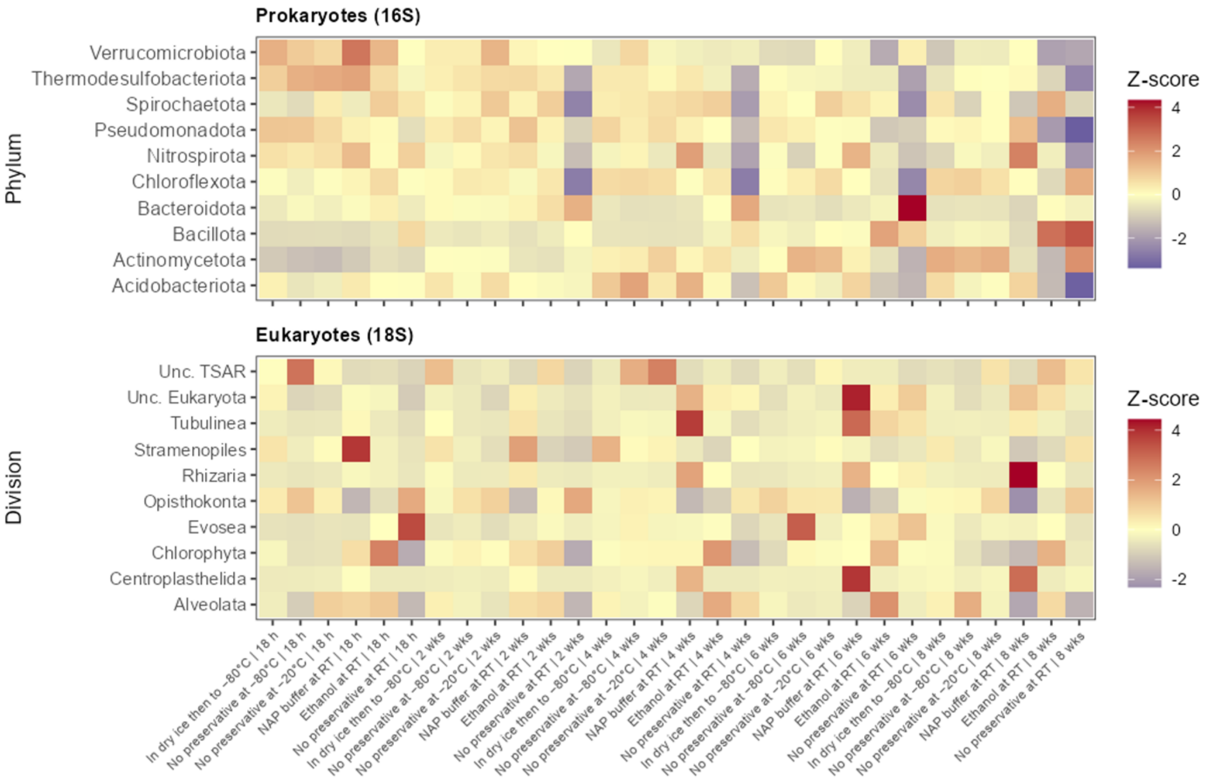


**Supplementary Figure S2.** Heatmaps showing the 10 most abundant bacterial phyla and eukaryotic divisions across preservation treatment × storage-duration combinations. Relative abundances were averaged within each treatment and time point and standardized within each taxon using z-score transformation.
